# Exploring the unmapped DNA and RNA reads in a songbird genome

**DOI:** 10.1101/371963

**Authors:** Veronika N. Laine, Toni I. Gossmann, Kees van Oers, Marcel E. Visser, Martien A.M. Groenen

## Abstract

**Background:** A widely used approach in next-generation sequencing projects is the alignment of reads to a reference genome. A significant percentage of reads, however, frequently remain unmapped despite improvements in the methods and hardware, which have enhanced the efficiency and accuracy of alignments. Usually unmapped reads are discarded from the analysis process, but significant biological information and insights can be uncovered from this data. We explored the unmapped DNA (normal and bisulfite treated) and RNA sequence reads of the great tit (*Parus major*) reference genome individual. From the unmapped reads we generated *de novo* assemblies. The generated sequence contigs were then aligned to the NCBI non-redundant nucleotide database using BLAST, identifying the closest known matching sequence.

**Results:** Many of the aligned contigs showed sequence similarity to sequences from different bird species and genes that were absent in the great tit reference assembly. Furthermore, there were also contigs that represented known *P. major* pathogenic species. Most interesting were several species of blood parasites such as *Plasmodium* and *Trypanosoma*.

**Conclusions:** Our analyses revealed that meaningful biological information can be found when further exploring unmapped reads. It is possible to discover sequences that are either absent or misassembled in the reference genome and sequences that indicate infection or sample contamination. In this study we also propose strategies to aid the capture and interpretation of this information from unmapped reads.

## Background

A vast amount of sequencing data is produced both at the DNA and RNA level. Often in next-generation sequencing projects, the starting point is the alignment of reads to a reference genome or transcriptome assembly, if such information is available. Despite improvements in alignment methods and hardware that have enhanced the efficiency and accuracy of alignments, a significant percentage of reads, frequently remains unmapped. Usually unmapped reads are discarded from the analysis process, but new biological information can be uncovered from this data. Such information can be e.g. pathogens, symbionts or sequences/genes missing in the reference genome [1–5].

Effort has been put into using already existing and/or creating new bioinformatics tools, especially for exploring pathogens in human sequence data [1, 6, 7]. In a study of the unmapped reads generated by the 1000 Genomes Project [8] biologically relevant information was identified from the reads that were non-human such as e.g. human papilloma virus [9]. In addition to known pathogens, also novel pathogens can be found (i.e. hitherto unknown pathogens or host-pathogen interactions). In a study of unmapped reads of the bovine reference individual many reads represented invertebrate species, some of which had an unknown link to bovine species [4]. These include parasitic infections but may also lead to the discovery of previously unknown symbiotic relationships. In a study of pea aphids (*Acyrthosiphon pisum*) that focused on symbionts, the symbiont sequences from the unmapped reads were most frequently shared between individuals adapted to the same host plant [3], indicating that these sequences may contribute to the divergence between host plant specialized biotypes.

One of the common findings in studies exploring unmapped reads is the incompleteness of the reference genomes especially at the gene annotation level. The so-called “missing genes” have been a problem especially in avian genomes. The recent sequencing and annotation of a large number of avian genomes [10, 11] as well as non-avian reptile genomes [12] made it possible to identify genomic features that are only found in birds, and that are linked with the evolutionary emergence of avian traits. However, one of the surprising findings of avian comparative studies is the loss of protein coding genes, as the total number of uniquely identified avian coding genes is considerably smaller than for other tetrapods [10, 13, 14]. In an analysis of 48 bird species, the total number of genes in avian genomes was estimated to be around 70% of those present in humans [10]. When investigating 60 avian genomes it was found that birds lack approximately 274 protein coding genes that are present in the genomes of most vertebrate lineages [14]. In another study it was highlighted that some of these 274 missing genes could be assembled from bird sequence data deposited at public databases ([15] but see also [16–18]). They suggested that the high GC-content of the missing genes could have caused problems in the PCR amplification in next-generation sequencing library preparation, as GC-rich genes are extremely hard to amplify [19].

A recent study of bird genomes and transcriptomes revealed that birds most likely do not contain fewer genes than mammals or non-avian reptiles [20]. These results indicate that the studies mentioned above have overlooked roughly 15% of the bird gene complements. They showed that there is a strong effect of local GC base composition, with genes with high GC-content being the most difficult to reconstruct consistently across different bird assemblies. However, they also were able to reconstruct missing genes with moderate or low GC-content, hinting that GC composition is not the only reason why so many bird genes have been overlooked so far. Because bird genomes are characterized by an extremely stable karyotype and recombination landscape, including GC biased gene conversion [21], it is very challenging to ultimately proof the absence of a particular gene within a genome. Hence, the question remains how many avian genes are truly missing from their genomes and how many are just not properly assembled and annotated.

The great tit (*Parus major*) is a well-known model species for ecological and evolutionary studies with several long-term study populations [22]. In addition to ecological and evolutionary studies, research of great tits and their pathogens has contributed to a vast knowledge on host-parasite coevolution [23]. Furthermore, many molecular datasets have been generated for this species, resulting in an extensive number of molecular tools [24–26]. However, although the genomic information for great tit is one of the most comprehensive among birds, the annotated genome still has some limitations such as the absence of some chromosomes such as micro-chromosomes that are missing in other birds as well, chromosome 16 and the sex chromosome W (as the reference bird was a male and thus was lacking the W chromosome). Especially chromosome 16 is known to be problematic to assemble in birds, since it contains the highly polymorphic MHC-gene complex region [27]. In addition, there are still regions in the great tit genome where no sequences have been assigned, most probably due to extreme GC-content and repetitive elements.

In a previous study by Santure et al. [28], the first great tit transcriptome was described using RNA extracted from ten birds and eight different tissues. When exploring the unmapped RNA reads, they found interesting signals from contaminants and pathogens. However, at that time the great tit reference genome was not yet available. Here, we explored the great tit unmapped reads further using an extensive dataset generated from the great tit reference individual, in order to flag problematic areas in the genome and identify pathogens and contaminants. For this, we used the unmapped reads of the DNA (normal and bisulfite treated) and RNA sequencing data from nine different tissues of the reference bird.

## Methods

### Sampling, extraction and sequencing

The workflow for this study has been outlined in Figure 1. We used DNA sequencing data of blood, bilsulfite treated DNA sequencing data of blood and brain and RNA sequencing data of eight tissues derived from the individual used to generate the great tit reference genome (BioSample: SAMN03083587) and submitted previously to NCBI SRA-database. Sample preparation, DNA and RNA extraction and sequencing for these tissues have been described previously[25]. In short, DNA was extracted from whole blood of the reference bird and sequenced with Illumina HiSeq 2000 at ~ 95X. The DNA sequencing data have been deposited in NCBI (SRX1539210, SRX1519144, SRX1517153, SRX1517152, and SRX1517034). Blood and brain DNA libraries were constructed according to the Epitect whole-genome bisulfite sequencing workflow (Illumina) and the whole-genome sequencing data were generated using the Illumina HiSeq 2,500 platform. The methylation data has been deposited in NCBI with accession numbers SRR2070790 and SRR2070791 for the brain and the blood, respectively.

**Figure 1.**
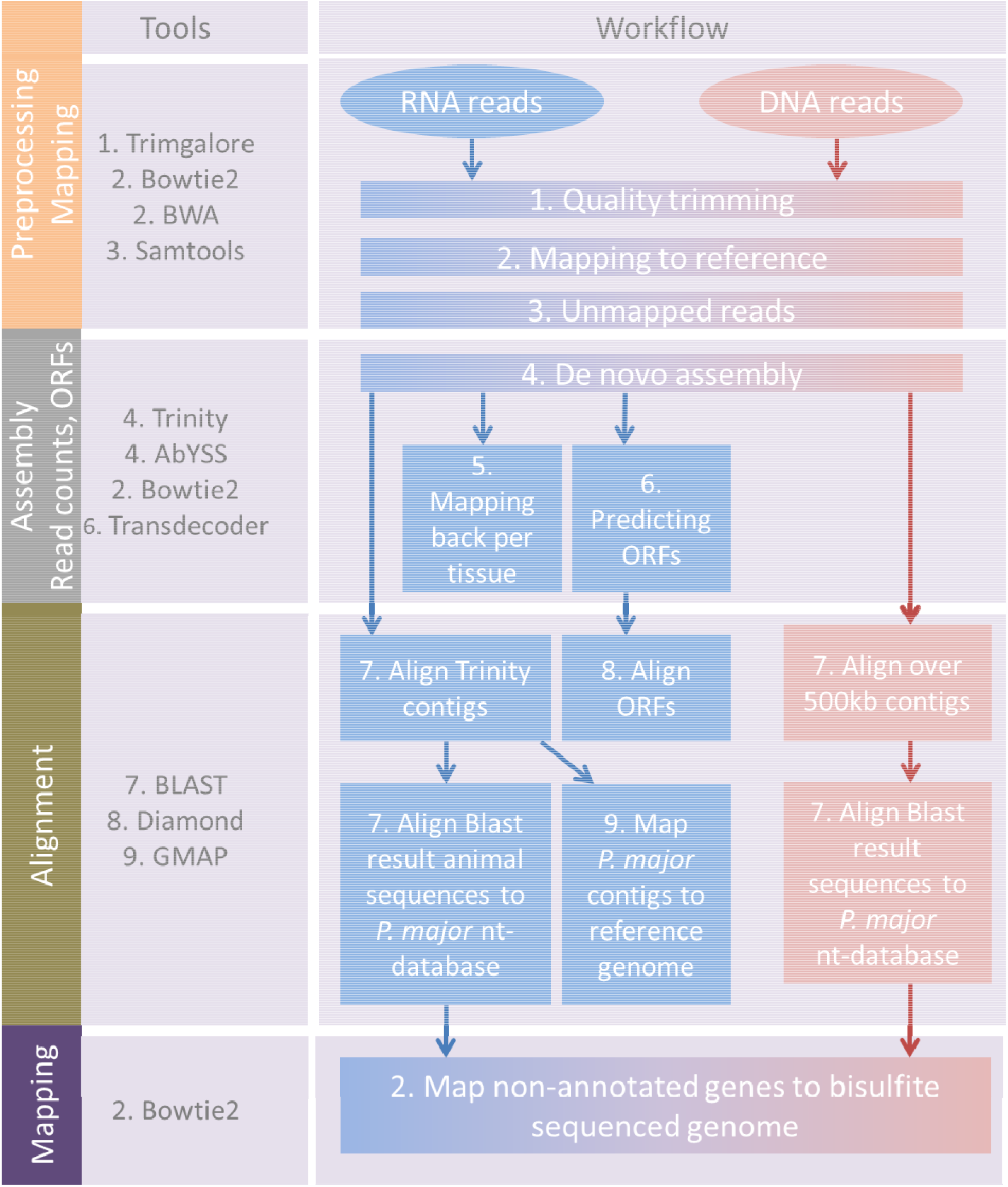
Schematic overview of the workflow used for the analysis of unmapped reads of DNA and RNA datasets.

RNA was extracted from eight tissues (bone marrow, homogenized half of the brain, breast filet, higher intestine, kidney, liver, lung and testis) from the reference bird and was then used to prepare tissue-specific tagged Illumina sequencing libraries. The tagged libraries were pooled and sequenced using five lanes on one flowcell of Illumina HiSeq 2000. This resulted in 100 bp paired-end unstranded RNA sequencing data. The number of reads per tissue ranged from 98 to 229 million with a total number of 1.25 billion paired-end reads. For the current study we also sequenced RNA isolated from whole blood of the reference bird where the majority of the RNA comes from the nucleated red blood cells. Blood RNA extraction was done with Direct-zol RNA miniPrep Plus kit (Zymo Research) with a modification in the start of the protocol. For the sample preparation we used 100 μl of blood in EDTA buffer mixed with 300 μl Trizol and shook for 5 minutes with vortex and by hand and then proceeded with the RNA Purification by following the protocol. The blood library was sequenced using a single lane on the Illumina HiSeq 2500.This resulted in 125 bp paired-end unstranded RNA sequencing data. The number of read pairs was 259 million. All the reads were checked for quality using FastQC [29] and low-quality sequences were trimmed with Trim Galore v. 0.4.4 [30] retaining unpaired reads resulting in a final number of 1,436,348,370 paired-end reads and 39,230,998 unpaired reads. The RNA sequencing data per individual tissue have been deposited in NCBI (GT_BoneMarrow SRS863935, GT_Brain SRS866013, GT_BreastFilet SRS86603, GT_HigherIntestine SRS866033, GT_Liver SRS866035, GT_Kidney SRS866036, GT_Lung SRS866044, GT_Testis SRS866048, GT_Blood SR[to be submitted]).

### Mapping and alignment of the DNA reads

After quality trimming with Trim Galore the reference bird DNA reads were aligned to the reference genome with BWA v.0.7.15 [31] using the default settings. The unmapped reads were obtained with Samtools [32] and subsequently assembled with AbYSS v. 1.3.7 [33] with k=20. Contigs larger than 500 bp, were aligned using Blastn against the BLAST nt-database, followed by aligning the resulting sequence hits (e-value < 1e-10) against only to *P. major* nt-database the same as for the RNA sequencing data described below.

### Assembly and mapping back of the unmapped RNA reads

The RNA reads from the nine different tissues were mapped against the great tit reference genome (NCBI *Parus major* genome version 1.1, GCA_001522545.2) using Bowtie2 [34]. We mapped paired and unpaired reads separately, using --local --very-sensitive-local options. Mapping success was compared with splice-aware mapper Hisat2 [35] which showed lower mapping percentages than Bowtie2, marking many great tit sequences as unmapped (Additional file 1: Table S1), which made us to use Bowtie2 mapping results in the end. The unmapped reads were obtained with Samtools and transformed to fastq –reads with Picard [36]. From the unmapped reads a *de novo* assembly was generated using Trinity [37]. In order to get read counts for every tissue separately, the unmapped reads were mapped back to the Trinity assembly using Bowtie2 and read counts were obtained with Samtools.

### Alignment of unmapped RNA-contigs to the BLAST database

The closest matching sequence to each Trinity contig was identified by alignment to the NCBI non-redundant nucleotide (nt) database using Blastn and keeping only the best hit for each contig provided that e-value was below 1e-10. Furthermore, we also predicted open-reading frames with Transdecoder [37] and used DIAMOND [38] to get the closest peptide match. Based on the taxid in the blast results, the contigs were divided into five groups: great tit transcripts, other bird transcripts, other animal transcripts, plants and other. The contigs that were identified to be great tit sequences in the BLAST analysis were mapped back to the great tit reference genome using GMAP [39] in order to get the exact genomic positions and mapping qualities. Using longer sequences such as assembled contigs instead of sequence reads might improve the mapping success to reference.

### Identifying the missing genes from the *P. major* annotation

From the BLAST results in total 4,784 nt-database sequences, classified as “other bird” and “other animal” species groups (e-value lower than 1e-10), were aligned to the nt-database using Blastn but restricting this search only to *P. major* sequences in order to see if there was a corresponding gene in the *P. major* annotation and thus avoiding gene naming differences between species. The corresponding Trinity contigs of the sequences that were not aligned to the *P. major* nt-database were treated as missing genes from the *P. major* annotation.

### Mapping of bisulfite sequenced genome

We extracted the unmapped reads from the bisulfite sequences (blood and brain tissues [25, 40] mapped to the reference genome with Bismark [41]. These unmapped bisulfite reads were assembled with AbYSS v. 1.3.7 with k=20. Contigs longer than 100 bp were aligned against the whole BLAST nt-database using Blastn, followed by aligning the resulting sequence hits (e-value < 1e-10) against only to the *P. major* nt-database in a similar way as with the DNA sequencing and RNA sequencing data described above.

During bisulfite sequencing, unmethylated cytosines are transformed into uracils, and hence the GC content of the reads will be lower in comparison to standard genome sequencing. We hence tested whether differences in GC-content between the two sequencing approaches have an effect on transcriptome mapping success. We used Bowtie 1.2.2 [42] to map bisulfite sequence reads and standard genome sequenced reads to our newly identified gene set from the RNA dataset and, as a comparison, to the latest release of the RefSeq annotated genome of *P. major* (release 1.1). We used bisulfite genome sequences from blood from the reference bird (BioSample: SAMN03781031). To obtain coverage distributions, we counted average per base coverage across transcripts using Samtools depth.

### Constructing the mitochondrial genome of the reference bird

The mitochondrial reads were extracted from DNA sequencing reads by aligning the reads to an already published *P. major* reference mitochondrial genome (NC_026293) with BWA. The aligned reads were then assembled using Geneious 9.1.8 [43]. For the annotation we used MITOS [44]. The newly constructed mitochondrial genome, the already existing reference mitochondrial genome and 123 sequences of NADH dehydrogenase subunit 2 (*ND2*) gene from both *Parus major* and *Parus minor* were used in a phylogenetic analysis. This was conducted with Geneious 9.1.8 and pairwise distances were estimated using the Tamura-Nei distance method and Neighbour-Joining was used to generate a phylogenetic tree.

## Results

### De novo assemblies of unmapped reads

Around 38.5 million DNA sequence reads, 3.62% of the total, remained unmapped after alignment to the reference *P. major* genome. These reads could be assembled into 1,064,033 contigs (N50 = 770bp) of which 1053 were larger than 500 kb. Additional assembly statistics are provided in Additional file 1: Table S2.

A total of 248 million RNA reads were unmapped (9% of the total). From the different tissues the intestine and bone marrow samples contained the largest fraction of unmapped reads (over 20% per tissue; Figure 2). A *de novo* assembly of these reads yielded altogether 136,122 contigs with an N50 of 1747 bp and a median contig length of 559 bp. From these contigs 80,435 open-reading frames could be identified. Additional assembly statistics are provided in Additional file 1: Table S3.

**Figure 2.**
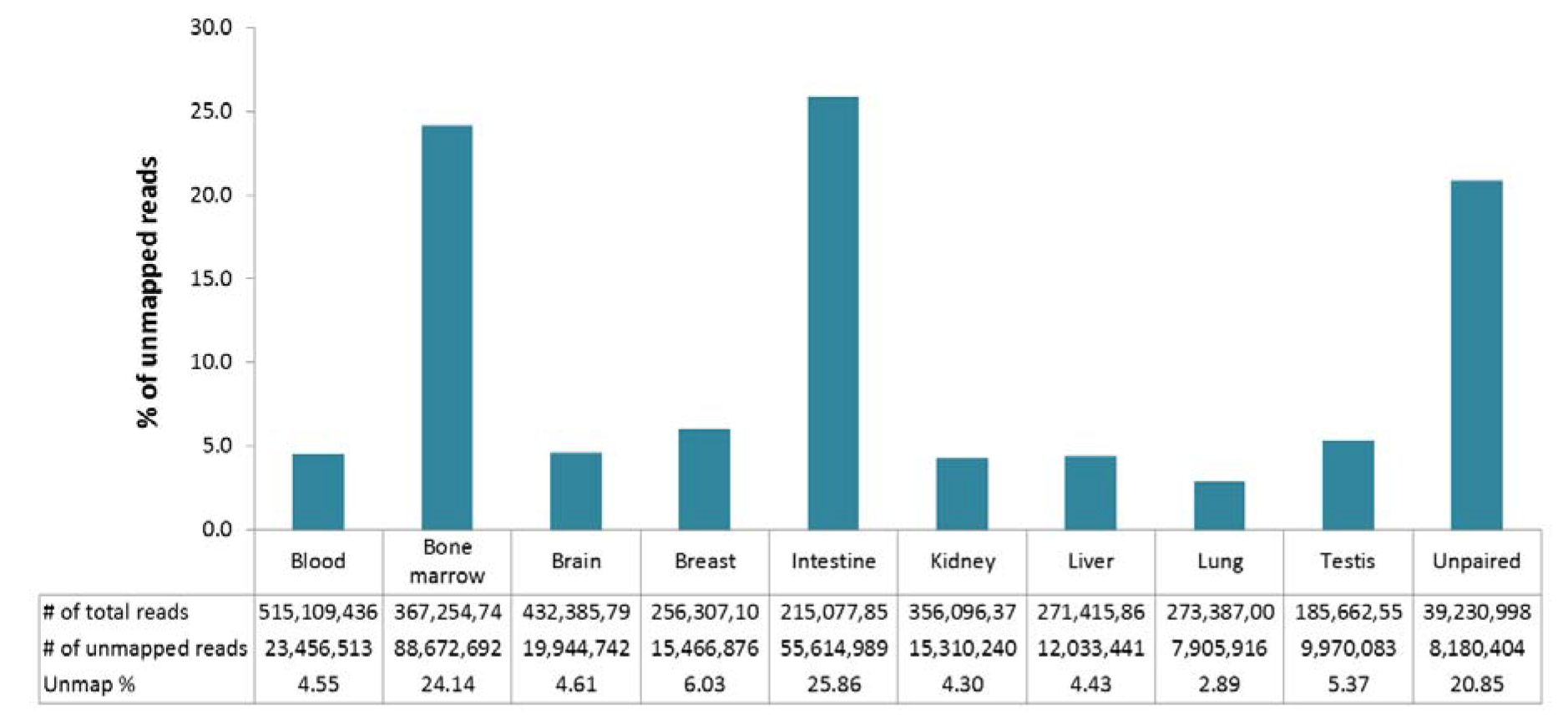
The percentage and number of the RNA reads that were unmapped to the reference genome in nine great tit tissues. The unpaired reads indicate the reads that were orphaned in the quality trimming of the RNA reads (all the tissues combined).

### Alignment of DNA contigs to BLAST database

From the assembled DNA contigs, 396 aligned against sequences from other birds in the BLAST run, with most of the contigs aligning to the ground tit (*Pseudopodoces humilis*) and blue tit (*Cyanistes caeruleus*)(Additional file 1: Table S4). Altogether 241 contigs could be aligned to a gene (actual or prediction), 154 of which were already identified in the great tit annotation. The GC-content of these great tit genes was 51.41% and gaps in the genome were found in five genes. The 87 remaining genes were missing from the great tit annotation. The GC-content of the contigs related to these genes was 51.94%.

### Alignment of RNA contigs to BLAST database

Altogether from the assembled Trinity contigs, 88,209 could be aligned to the non-redundant nucleotide (nt) database, of which 85,771 had an e-value lower than 1e-10 (Table 1). Among the open-reading frame peptides, 65,942 had a significant alignment to the peptide database (Additional file 1: Table S5).

**Table 1.** Summary of the BLAST alignments of unmapped RNA contigs with e-value less than 1e-10.

| Group | Number of alignments | Median identity (%) | Maximum identity (%) | Median qcovs (%) | Maximum qcovs (%) | Median E-value |
| --- | --- | --- | --- | --- | --- | --- |
| *Parus major* | 217 | 97.07 | 100 | 100 | 100 | 4.27E-113 |
| Aves | 20,883 | 96.35 | 100 | 52 | 100 | 1.35E-92 |
| Other animals | 569 | 91.01 | 100 | 82 | 100 | 2.78E-82 |
| Plants | 62,019 | 100 | 100 | 99 | 100 | 0 |
| Other | 2084 | 100 | 100 | 100 | 100 | 2.19E-125 |

The most common alignments were to plant and to other bird sequences. The plant sequences were mostly derived from *Arabidopsis* (Additional file 1: Table S6). The vast majority of these *Arabidopsis* reads was present among the unmapped reads from bone marrow suggesting contamination of this library with plant derived sequences (Table 2). Contigs related to other bird species were equally distributed among tissues except for intestine that had five times more reads aligned (Table 2). The reads from the intestine were mostly aligning to trypsin related genes (Additional file 1: Table S7). Most of the *P. major* contigs originated from the breast filet sample and the unpaired dataset. These contigs mostly aligned to great tit mitochondrial sequences in the nt-database and when aligning these contigs against *P. major* genome, they mapped back with low MAPQ –values (0-3) especially to the mitochondrial genome (Additional file 1: Table S6 and S7). The GC-content of the non-mitochondrial contigs was 52.7% and 26% of the contigs had repeats in them. In the “other” group, bacterial reads were the most common type in almost all of the tissues, except in blood where fungi and other eukaryotes were the most common species groups (Table 2, Figure 3). The reason for the high number of fungal and other eukaryotic sequences in blood was due to ribosomal RNA sequences (Additional file 1: Table S7). The most prevalent organisms in the “other” group were *Plasmodium relictum* and *Caldibacillus debilis* (Additional file 1: Table S6 and S7). When looking at the *Caldibacillus* record in Genbank (MF169985.1) more closely we noticed it had contamination and was being removed from the nucleotide database. Thus, we re-did the alignment for these contigs and the results showed that one of the contigs was aligned with *Actinomyces succiniciruminis* and rest of the contigs (6) aligned to *Culicoides sonorensis* but closer look at the sequences revealed them being PhiX control reads (used as a quality and calibration control for sequencing runs) indicating PhiX contamination in many of the NCBI submitted sequences.

**Figure 3.**
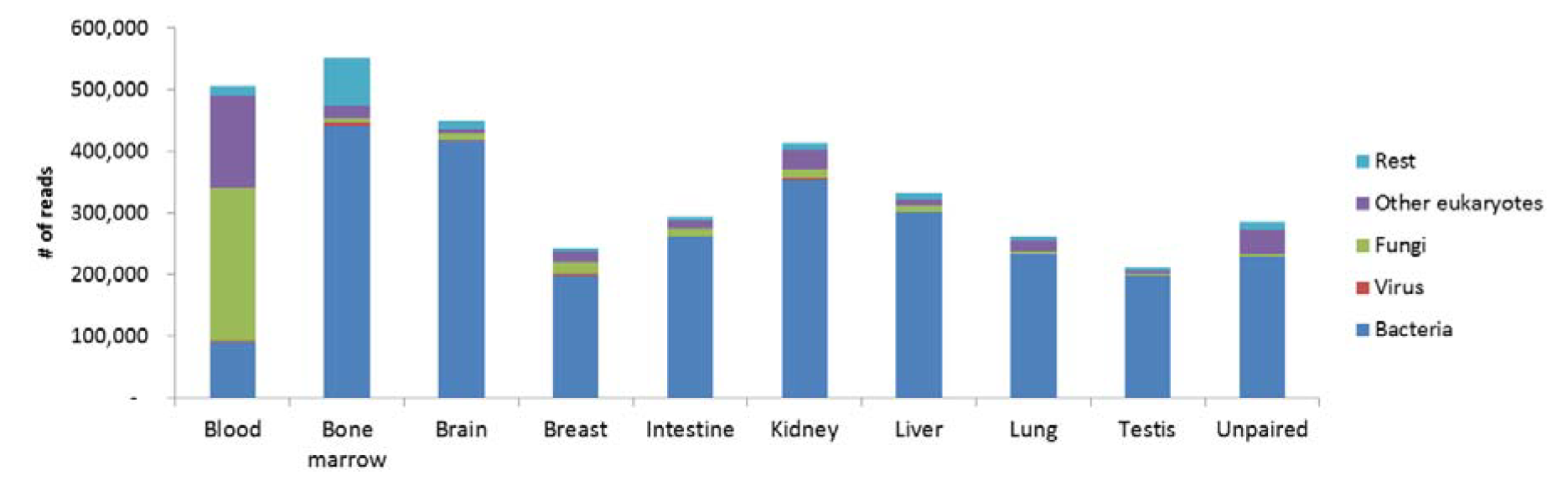
Closer examination of the “other” –group from Table 2 per tissue type.

**Table 2.** RNA read counts per tissue type and per major BLAST alignment result group. The unpaired reads indicate the reads that were orphaned in the quality trimming of the RNA reads (all the tissues combined).

| Group | Blood | Bone marrow | Brain | Breast | Intestine | Kidney | Liver | Lung | Testis | Unpaired |
| --- | --- | --- | --- | --- | --- | --- | --- | --- | --- | --- |
| *Parus major* | 4,295 | 19,657 | 33,516 | 300,763 | 55,999 | 72,625 | 41,800 | 20,834 | 13,693 | 935,271 |
| Aves | 10,796,873 | 6,087,566 | 9,646,728 | 8,395,675 | 48,812,307 | 7,570,425 | 5,637,931 | 3,758,819 | 5,039,638 | 9,193,238 |
| Other animals | 2,109,001 | 435,891 | 581,626 | 572,849 | 303,246 | 429,018 | 245,629 | 177,476 | 128,091 | 428,660 |
| Plants | 1,949,545 | 75,408,648 | 399,632 | 409,233 | 325,667 | 559,343 | 344,760 | 244,482 | 182,009 | 1,982,031 |
| Other | 506,723 | 552,931 | 449,697 | 244,110 | 295,119 | 413,895 | 332,886 | 262,519 | 212,792 | 285,692 |
| Sum | 14,493,076 | 82,422,401 | 11,016,164 | 9,732,532 | 49,708,365 | 8,918,388 | 6,534,019 | 4,412,787 | 5,548,801 | 12,779,187 |

### Identifying the missing genes from the *P. major* annotation by using RNA sequencing data

From the 4,784 nt-database sequences that were related to other bird and animal species, 2,625 did not align to great tit when restricting the alignment only to the *P. major* nt-database. These corresponded to 12,759 Trinity contigs and the average GC-content of these contigs was 60.02%. From this we could identify 1,822 individual genes (1,110 gene predictions and 712 actual genes) that were missing in the great tit annotation for genome build 1.01 but that are found in the unmapped reads (Additional file 1: Table S8). Corresponding genes to *P. major* annotation could be found for 1,931 nt-database sequences and the rest of the sequences (227) did not align, since these were sequences from complete chromosomes or un-assigned scaffolds (Additional file 1: Table S6).

From these genes gaps were found in 68 genes and the GC-content was slightly higher (GC% 54.51) than the average for the whole genome (GC% is 41.52).

### Bisulfite sequence genome information

A total of 217 and 341 million bisulfite reads from blood and brain, respectively (41.6% and 54.1% of total reads), were unmapped. The unmapped reads from the bisulfite data could be assembled into 145 million contigs (with N50 of 836 bp) in blood, of which 929 were larger than 100 kb and into 96 million contigs (with N50 of 1 259 bp) in brain, of which 1,956 were larger than 100 kb (Additional file 1: Table S9). For both tissues, the unmapped reads aligned mostly against *E. coli* in the BLAST run (Additional file 1: Table S10). Altogether 59 and 226 contigs in blood and brain, respectively, could be aligned to a gene (actual or predicted). Of these genes, 12 from blood and 59 from brain were not found in the great tit genome, most of them being prediction genes from blue tit (*Cyanistes caeruleus*) and ground tit (*Pseudopodoces humilis*).

If extreme heterogeneity in local GC composition is causing sequencing issues and hence unequal coverage, the ability to reconstruct genes from these regions may be affected. We would expect that bisulfite sequenced genome reads should show a different mapping success in comparison to standard sequenced genomes. Indeed if we map the reads to the RefSeq transcripts, i.e. currently annotated known great tit genes, the mapping behaviour is remarkably different between classic whole genome reads and bilsulfite treated genome reads (Figure 4 A and C). To our surprise, for our newly generated gene datasets, we do not see such an effect (Figure 4 B and D).

**Figure 4.**
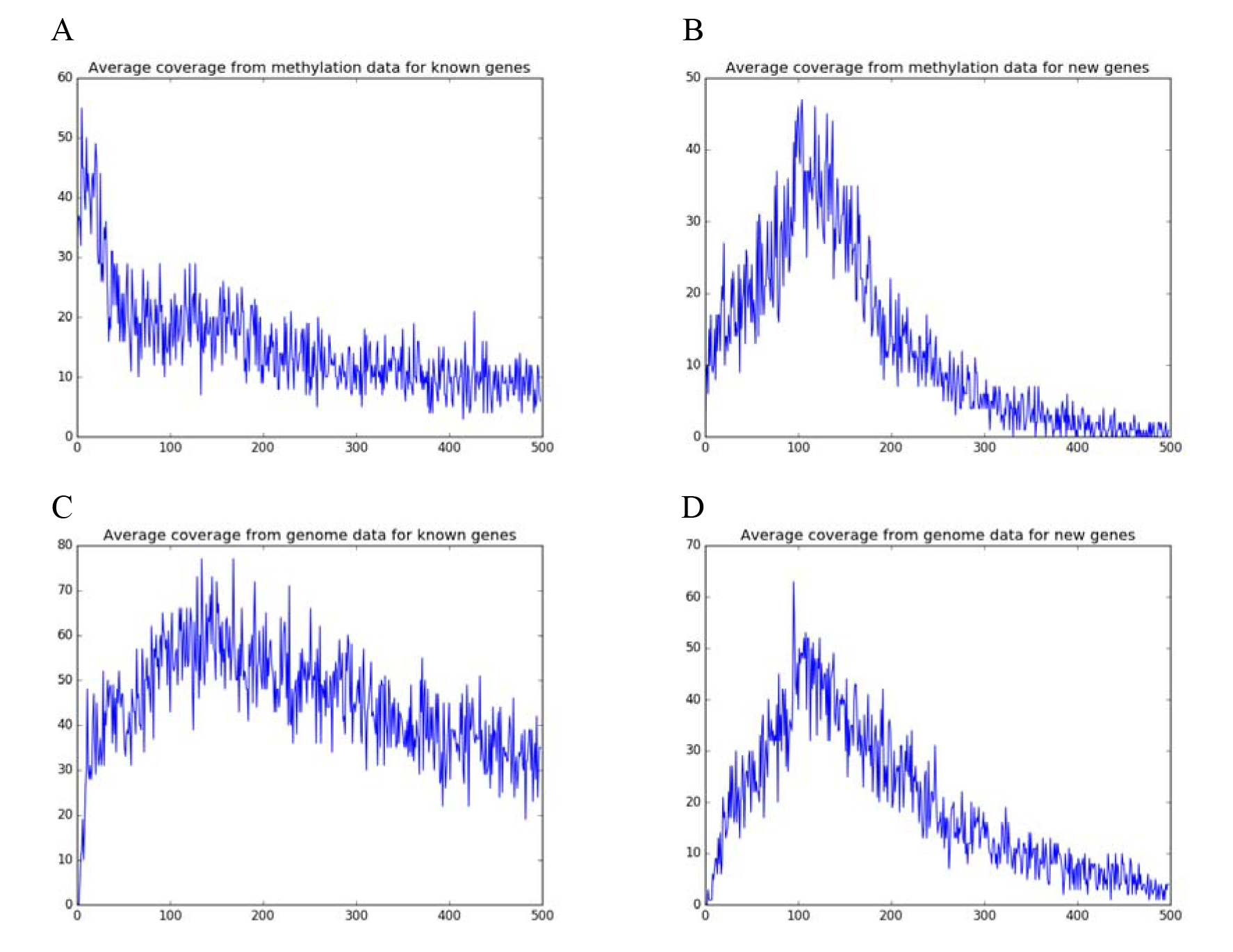
Histograms of average per base coverage of unconverted whole genome sequence data and bisulfite converted whole genome data onto the Refseq transcripts (“known genes” - A,C) and the newly identified genes in this study (B,D).

### The mitochondrial assembly

From the unmapped DNA reads we could assemble the complete mitochondrial genome for the reference genome bird. The complete mitochondrial genome of *P. major* is 16,777 bp long [to be submitted to NCBI] and contains 37 genes (Additional file 1: Table S11 and Figure S1). In the phylogenetic analysis the newly constructed mitochondrial assembly grouped with the *P. major* samples and the Chinese *P. major* reference mitochondrial genome grouped with the *P. minor* samples (Figure S2).

## Discussion

Our analyses revealed that meaningful biological information can be obtained from analyzing unmapped reads. We discovered sequences that were either absent or misassembled in the reference genome and detected sequences that indicated infection or sample contamination. Furthermore, investigating unmapped reads helped us to discover species relationships, especially pathogenic, for the great tit.

### Contamination

In our RNA sequencing dataset a large amount of plant sequences was detected in the unmapped reads, mostly in the bone marrow sample. In the bone marrow over 90% of the unmapped reads were from *Arabidopsis* suggesting that there has been a contamination during the library preparation or sequencing. Another contamination signal was found from the bisulfite data in both tissues with a high number of *E. coli* sequences. Contamination can cause problems in next-generation sequencing projects especially if the contaminant species is related to the study species. Tae et al. [9] found specific contaminants that are linked to certain sequencing centers and they also found reads that were falsely classified as contamination, because of the high similarity of human sequences to sequences in non-human genome assemblies such as mouse. Contaminant-derived reads that are mapped to the genome can give false information, or if used in a genome assembly, can cause misassembly of sequence contigs leading to erroneous conclusions and incorrect annotation of genes that are absent in the study species. The *Caldibacillus* incidence in our RNA data where the actual hit was PhiX, shows that published contaminated sequences deposited in sequence archives can have an impact on other studies and affect the interpretation of the results if the PhiX reads are not filtered well enough [45].

### Pathogens

Sequences representing possible pathogenic invertebrates, viruses or prokaryotes, were identified in all of the tissue types in the RNA sequencing dataset. Out of the 2084 Trinity contigs that were identified as “others” and possible pathogenic, 1491 were derived from *Plasmodium relictum*, one of the causes of avian malaria, which is known to infect a large number of bird species and targeting especially great tits [46]. *Plasmodium* was very abundant in our samples suggesting that the reference bird was a carrier for this pathogen. Although the reference bird was captive bred, it resided in an open-aviary so there is a possible route to become infected through mosquitos, fleas and mites. Moreover, avian malaria infections might originate from early infections during the nestling phase, since there they are more susceptible to the vectors[47]. Captive bred individuals have experienced a 10-day period in a natural nest in a wild population, and therefore, these infections could likely originate from that phase [48]. The observed *Trypanosoma* reads were mostly found in blood and bone marrow samples, the typical tissues this parasite is found in birds [49]. Birds in the aviary facilities experienced a severe *Trypanosoma* outbreak at the time the reference bird’s mother was housed there, indicating that the vectors distributing these parasites are present when keeping birds in captivity. There were also pathogens not previously linked to birds, which might suggest these pathogens infect a larger group of species or have a related species that infect birds but that has not been sequenced before. If there is a sufficient number of pathogenic reads, it is possible to measure the expression of genes from these pathogens and link it to certain host tissues if several tissues have been sequenced. Furthermore, knowing the infection status of studied individuals is important. Especially in a study where different groups of individuals are compared it is important to know if some of the individuals are infected as this can affect the results for example of a differential expression study. For an example an undetected malaria infection can have a huge impact on behavioural studies in great tits as it has been shown that differences in malaria load can affect the personality of the birds [50]. Next generation sequencing methods have not been fully utilized in the host-parasite studies of great tits, which might prove to be a successful avenue to explore in addition to visual inspections and PCR-related methods.

Another possibility for finding these non-vertebrate sequences in unmapped RNA sequencing reads is the actual integration of these sequences into the great tit genome. Horizontal gene transfer can maintain pre-existing functions or can provide new functionality for the recipients, which can lead to e.g. adaptation [51]. Horizontal gene transfer from prokaryotes to eukaryotes has been reported to occur in many animal species [51] but studying them needs a careful planning due to the complexity of the genomes [52, 53]. Illustrative, especially in human genome studies, are several misattributions of horizontal gene transfer events [53, 54]. Integration of foreign DNA released by dead cells into healthy host cells is also possible [55, 56]. Nothing is known about horizontal gene transfer in the great tit and in our BLAST results we did not find any contig that was partially mapped to bird and partially mapped to a non-vertebrate species. Horizontal gene transfer remains an interesting avenue to follow in more detail in the future.

### Flagging problem areas in the genome assembly

The BLAST sequence alignments showed that many of the assembled transcripts represented sequences from other bird species and that many of them were derived from genes that were already annotated in the great tit genome. Close inspection of these genes and their sequences revealed gaps in the genome sequences for some of them and also showed that the GC-content of these sequences was higher than the average GC-content of the genome. There were also contigs that were aligned to *P. major* sequences in the BLAST analysis. One explanation is, that we were unable to map these RNA sequences to the reference genome due to the mapping tool used, in this case Bowtie2 since this is not a splice-aware mapper that might discard reads that span over two exons. However, we also tested Hisat2 which takes splicing into account but this program discarded even more great tit related reads suggesting the problem might lie somewhere else. The majority of the unmapped great tit specific reads had their origin in the mitochondria. The presence of mitochondrial sequences in the unmapped reads can be explained by the fact that the reference mitochondrial sequence in GenBank does not come from the reference bird, but from a *P. major* sample collected from China [57]. When comparing our mitochondrial assembly to this Chinese one and also adding *ND2* gene GenBank sequences from both *P. major* and *P. minor* we could show that the Chinese mitochondrial genome actually comes from a *P. minor* individual. The rest of the great tit genes that had unmapped reads, were all gene predictions and the contigs linking to these genes had high repeat and GC-content hinting that these genes are problematic in general.

### GC content features of missing genes identified and TRY1 gene expression

*P. major* annotation release 1.01 consists of 18,744 annotated genes and pseudogenes. In our study we could find genes that were missing from the *P. major* annotation. The GC-content for these contigs was relatively high which could partly explain why these genes are hard to assemble. To investigate this further, we used blood whole genome sequence data with and without bisulfite pre-treatment (which reduces GC-content) and mapped it onto the *P. major* annotated genes and our newly identified gene set. We identified mapping differences between bisulfite treated and untreated DNA mapped to the known gene set, but not in our newly identified gene set. This is surprising, as these genes are GC rich and should therefore suffer more from sequencing issues in high GC content reads. One possible explanation is that these regions are highly methylated or contain fewer CpG regions. Indeed, gene body methylation is generally relatively high, in particular for lowly expressed genes [40]. We also observe a clear peak at around 100x for both sequencing methods for our newly identified gene set, suggesting that these genes might occur at several genomic locations (e.g. paralogs or pseudogenes). We hence conclude that our newly identified genes are not strongly affected by GC compositional sequencing effects, and that structural variation and context may play a more pronounced role why those genes have not been identified previously.

A prominent newly identified great tit gene was digestion related gene Trypsin I-P1 (*TRY1* / *PRSS1*), which was highly expressed in the reference bird and thus creating the tall peak in the intestinal tissue sample (Figure 2). This gene has been annotated in other birds but not in *P.major*. Hence, the used annotation can constrain especially gene expression studies if the unmapped reads are discarded without proper inspection. Long-read sequencing is often used to improve the reference genome as this can overcome the repeat area and GC-issues [58].

## Conclusions

We have shown that it is possible to find many sequences of interest from the reads that are not aligned to a reference assembly. These unmapped reads often provide biologically significant information such as identity and quantity of pathogenic organisms, possible contaminations and genes that are either partially or completely missing from the reference assembly. We also proposed strategies to aid the capture and interpretation of this information in great tit using unmapped reads. The composition of unmapped reads can be used in main research pipelines as covariate or phenotype. Especially in RNA studies, looking also at the gene expression of the missing genes can be beneficial. On its own, unmapped read research will also expand our knowledge of the extent of pathogens and symbionts. After all, a complex eukaryotic species, such as the great tit, is in fact a metagenome over time (horizontal gene transfer) and space (pathogens and microbiomes). We suggest when analysing NGS sequence data, especially from non-model organisms, to include reference databases from related species to avoid annotation biases and take particular care to distinguish contaminants from true biological meaningful signals.

## List of abbreviations

nt: non-redundant nucleotide database

## Ethics statement

Sampling of the reference bird was done under protocol number CTE 07.05 Adendum I, from the Animal Experiment Committee from the Royal Netherlands Academy of Sciences (DEC-KNAW) to KvO.

## Consent for publication

Not applicable.

## Availability of data and materials

The data set supporting the results of this article is available in the SRA repository, SRA accessions [to be submitted].

## Additional Files

TableS1. Number of RNA reads before and after trimming and mapping success of Bowtie2 and Hisat2.

TableS2. Summary statistics from the de novo assembly of unmapped reads from DNA sequencing using AbYSS.

TableS3. Summary statistics from the de novo assembly of unmapped reads from RNA sequencing using Trinity and all tissues combined.

TableS4. Summary of the significant alignments of de novo assembled contigs from DNA unmapped reads to the nt-database. Identity is the percentage of identical matches; query coverage is the query coverage per subject.

TableS5. Summary of the significant alignments of ORFs from RNA unmapped reads to the nr - database. Identity is the percentage of identical matches; mismatch is number of mismatches.

TableS6. Summary of the significant alignments of de novo assembled contigs from RNA unmapped reads to the nt-database. Identity is the percentage of identical matches; query coverage is the query coverage per subject; MAPQ is the mapping quality value from the mapping back to the *P. major* genome with GMAP.

TableS7. Read count per tissue type of the RNA dataset.

TableS8. List of newly discovered genes in the RNA dataset

TableS9. Summary statistics from the de novo assembly of unmapped reads from bisulfite sequencing using ABySS.

TableS10. Summary of the significant alignments of de novo assembled contigs from bisulfite treated unmapped reads to the nt-database. Identity is the percentage of identical matches; query coverage is the query coverage per subject.

TableS11. Annotation of the newly assembled *P. major* mitochondria. Score is the e-values for ncRNA and quality values for protein coding gene predictions.

FigureS1. Newly constructed and annotated mitochondria of the reference bird.

FigureS2. Neighbour-Joining phylogenetic tree between the newly constructed mitochondria (blue), the already existing reference mitochondria (green) and 123 sequences of NADH dehydrogenase subunit 2 (*ND2*) gene from both *Parus major* and *Parus minor*.

## Competing interests

Not applicable.

## Funding

VNL was funded by ERC Advanced Grant (339092 – E-Response) – awarded to MEV. TIG was funded by Leverhulme Early Career Fellowship Grant (ECF-2015-453) and a NERC grant (NE/N013832/1).

## Authors’ contributions

VNL designed the research, built the analysis pipeline and analysed the unmapped reads. TIG and VNL analysed the methylation data. VNL wrote the manuscript and TIG, KvO, MEV and MAMG edited the manuscript. All authors read the final manuscript.

## Acknowledgements

We thank Christa Mateman and Agata Pijl for lab assistance.

